# BiofilmQ, a software tool for quantitative image analysis of microbial biofilm communities

**DOI:** 10.1101/735423

**Authors:** Raimo Hartmann, Hannah Jeckel, Eric Jelli, Praveen K. Singh, Sanika Vaidya, Miriam Bayer, Lucia Vidakovic, Francisco Díaz-Pascual, Jiunn C.N. Fong, Anna Dragoš, Olga Besharova, Carey D. Nadell, Victor Sourjik, Ákos T. Kovács, Fitnat H. Yildiz, Knut Drescher

## Abstract

Biofilms are now considered to be the most abundant form of microbial life on Earth, playing critical roles in biogeochemical cycles, agriculture, and health care. Phenotypic and genotypic variations in biofilms generally occur in three-dimensional space and time, and biofilms are therefore often investigated using microscopy. However, the quantitative analysis of microscopy images presents a key obstacle in phenotyping biofilm communities and single-cell heterogeneity inside biofilms. Here, we present BiofilmQ, a comprehensive image cytometry software tool for the automated high-throughput quantification and visualization of 3D and 2D community properties in space and time. Using BiofilmQ does not require prior knowledge of programming or image processing and provides a user-friendly graphical user interface, resulting in editable publication-quality figures. BiofilmQ is designed for handling fluorescence images of any spatially structured microbial community and growth geometry, including microscopic, mesoscopic, macroscopic colonies and aggregates, as well as bacterial biofilms in the context of eukaryotic hosts.

## Main Text

Microbial biofilm communities shape the Earth by contributing to the biogeochemical cycles in soil, sediments, oceans, and the plant microbiota^1^. Microbial communities are also an integral part of human health, due to their functions in the intestinal and oral microbiota, as well as in infections, where cells that are bound in biofilms can display a 1000-fold higher tolerance to antibiotics than planktonic cells^2^. Biofilms are generally three-dimensional (3D) communities that display spatial gradients of nutrients and many other diffusible molecular compounds, as well as spatiotemporal variation in species composition and cellular differentiation^3,4^. Due to the spatial heterogeneity of phenotypes and genotypes inside biofilms, studies of biofilms often rely on 3D fluorescence imaging, *e.g*. using confocal microscopy. For biofilm phenotyping, and for characterizing phenotypes of particular cells within biofilms, it is critical to be able to perform image-based quantitative measurements of fluorescent reporters and structural features for particular regions inside the biofilm.

Extracting the desirable information from 3D images relies on non-trivial automated image analysis. The most widely-used tool for biofilm image analysis in the literature is COMSTAT^5^, which was released almost 20 years ago and provided one of the first tools to objectively determine differences in biofilm morphology. More recent software tools^6–8^ were released 10 years ago and include a refined threshold-based detection of biofilm biomass, the quantification of more fluorescence channels, and correlation functions. These tools have been tremendously important for biofilm research. However, since their release, there has been an ongoing revolution in image analysis capabilities, and in how quantitative information from biological images is extracted and presented^9^. The design of biofilm research projects and the discovery of new biofilm behaviours are currently limited by the lack of modern cytometry software tools that allow researchers to measure a comprehensive set of spatially and temporally resolved structural parameters and fluorescent reporters inside biofilms. New methods for the spatial and temporal analysis of biofilm structure, fluorescent reporters, and cytometry are therefore paramount to the field.

Inspired by the powerful and user-friendly tools for bacterial cell biology^10–13^, we have developed new image analysis algorithms for microbial communities, which we bundled in the form of a user-friendly software tool termed BiofilmQ (https://drescherlab.org/data/biofilmQ). This tool provides a graphical user interface that requires no programming or prior knowledge of image analysis; extensive documentation and video tutorials guide users through each step in the image analysis and data visualization pipeline.

BiofilmQ relies on standard fluorescence microscopy image data that does not need to resolve all individual cells. For 2D imaging, most fluorescence microscopy techniques should result in suitable images for BiofilmQ, while for 3D imaging we recommend using confocal or light sheet fluorescence microscopy to provide an optical resolution in the 3^rd^ dimension that is appropriate for the particular project under investigation. The core concept of the BiofilmQ 3D image analysis (Fig. 1a) is to first perform a 3D segmentation of the biofilm structure based on one of the fluorescence channels, to detect the 3D biofilm volume location, followed by the dissection of this 3D biofilm volume into a cubical grid, whose cubes have a user-defined size that can be chosen to be approximately equal to the cell volume. For further analysis, the individual cubes of this biofilm are then treated as pseudo-cells, for which data can be extracted as if they were single cells. This results in flow-cytometry-like data, but with the additional component of spatial features, without requiring optical single-cell resolution. For each cube, BiofilmQ quantifies as many fluorescence, structural, and textural parameters as desired, in addition to the cube’s spatial context (Fig. 1b). Independently from the pseudo-cell cubes, BiofilmQ computes global morphological, structural, textural, and fluorescence properties of the whole 3D biofilm volume (Fig. 1c). BiofilmQ therefore provides local and global quantifications of many biofilm properties. For the special case of 2D image data input (*e.g*. from epi-fluorescence, confocal, or any other fluorescence microscopy technique), the biofilm biomass area is segmented into small squares, analogous to the cubes in 3D images, followed by similar analysis steps to the 3D datasets. BiofilmQ thus allows biofilm cytometry and high-throughput phenotyping of biofilms, providing many spatially-resolved structural, textural, and fluorescence readouts that were previously unavailable.

**Figure 1:**
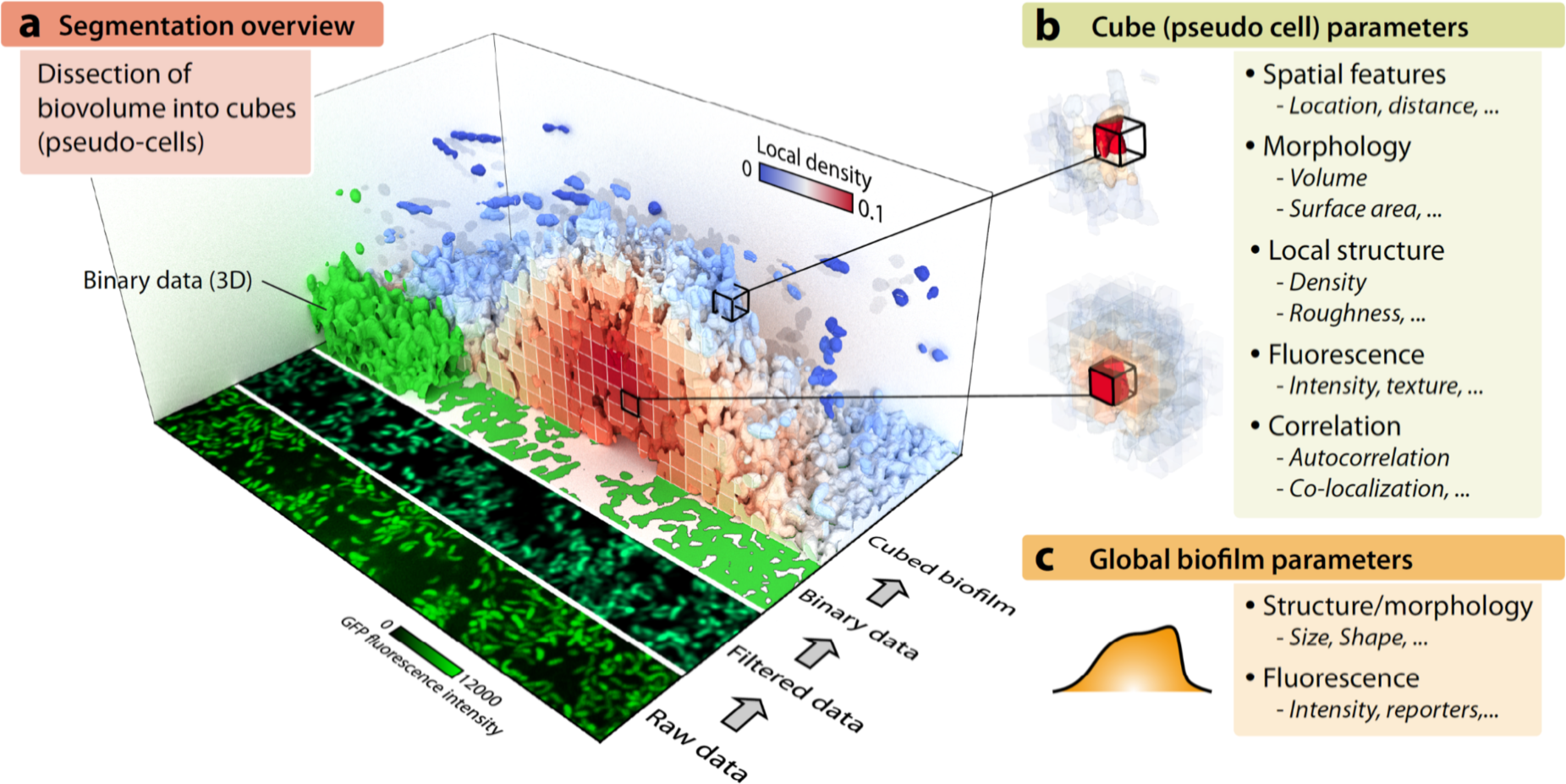
Overview of the image processing and parameter quantification concept of BiofilmQ. **a**, For a 3D biofilm image, the BiofilmQ image processing pipeline is illustrated: The original raw fluorescence image is filtered and thresholded to obtain a binary representation of the biofilm. This 3D binary data is then dissected into cubes (pseudo-cells) with user-defined size, to quantify the spatial distribution of structural, textural, and fluorescence properties. Here, each cube in the biofilm is coloured according to the local biomass density, which is one of the cube properties that can be extracted. **b**, Many parameters can be quantified for each pseudo-cell cube. **c**, Additionally, several global biofilm parameters are extracted.

Images of biofilms in any growth geometry and any species can be analysed with BiofilmQ (Fig. 2a) – the only requirement for the BiofilmQ analysis is that the 3D (or 2D) image dataset contains at least one fluorescence channel on which the community segmentation is based. Many commonly used microscopy image formats are supported in BiofilmQ (Fig. 2b), based on the Bio-Formats toolbox^14^.

**Figure 2:**
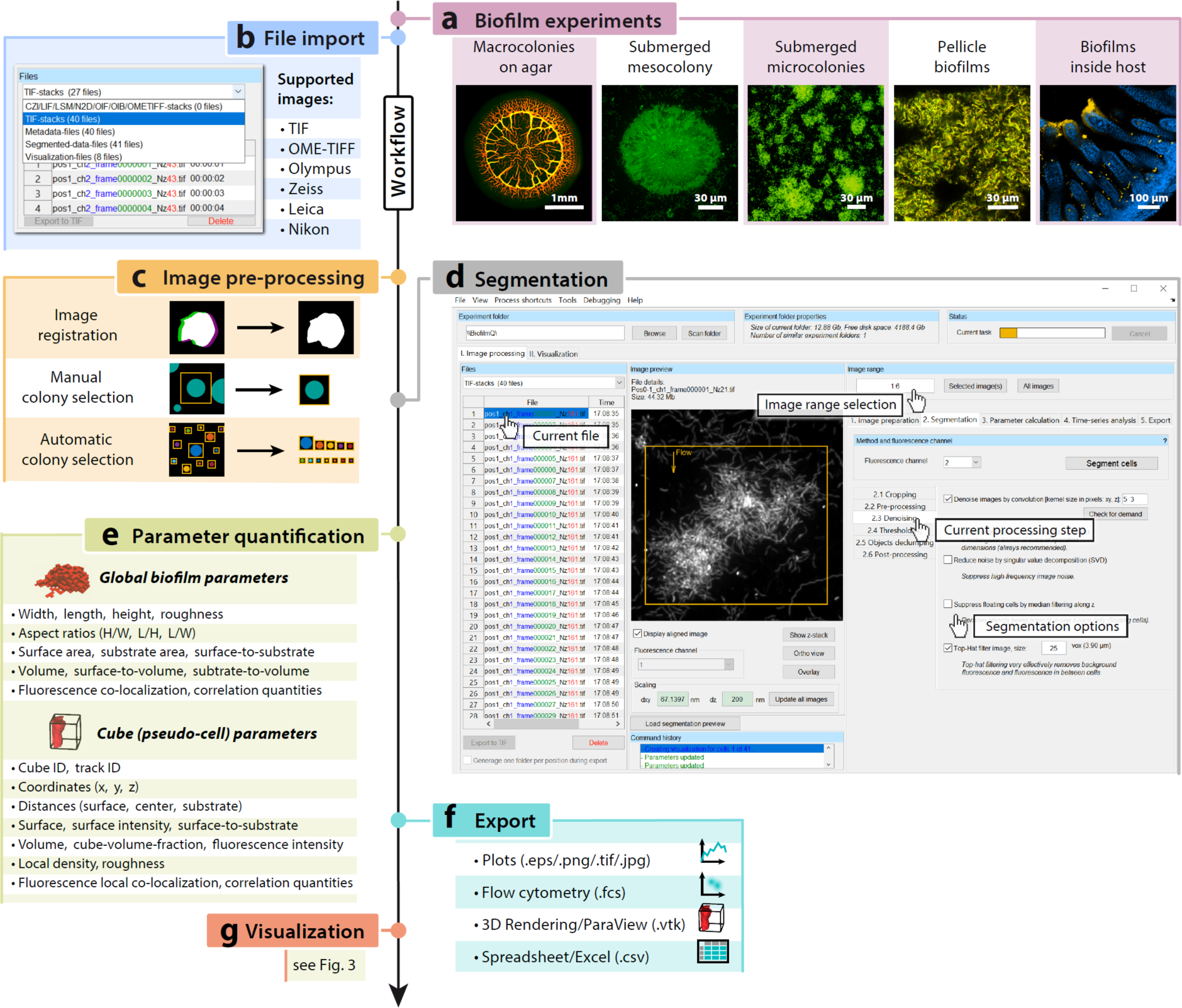
Workflow of the BiofilmQ user interface for image processing and analysis. **a**, Examples of biofilm growth configurations and image categories that can be analysed with BiofilmQ. **b**, First, images have to be imported; a wide range of formats are supported, including 3D image formats, and TIF image sequences. **c**, Next, optional pre-processing steps including image time series registration, filtering for noise reduction, and colony separation can be performed. **d**, During image segmentation, biomass is distinguished from background, which can be performed automatically using different thresholding algorithms, or semi-manually. After the segmentation of the biofilm volume, the biofilm is sliced into cubes of a user-defined size. **e**, The segmentation results can be used to extract quantitative information *via* the parameter calculation for the biofilm as a whole, and for each cube in the biofilm. **f**, Parameter quantifications and biofilm structural analysis can be exported either as spreadsheets, flow-cytometry data format, graphs, or as input data for a 3D rendering. **g**, The extensive visualization capabilities of BiofilmQ for the quantified parameters are described in Fig. 3.

An important first step in the quantification of biofilm properties is the segmentation of the biofilm biomass location (Fig. 2d). Biofilm segmentation algorithms have received significant attention in the literature^15–21^. BiofilmQ therefore offers different segmentation workflows: either semi-manual segmentation supported by immediate visual feedback, or *via* automatic segmentation algorithms, such as Otsu, Ridler-Calvard, robust background, or maximum correlation thresholding. Several optional pre-processing steps (Fig. 2c) may precede the threshold calculation to improve segmentation results.

Following the segmentation of the whole biofilm volume, the biofilm is dissected into cubes of equal size, which are treated as pseudo-cells for the local parameter calculation (Fig. 1c). This approach permits the spatially-resolved quantification of parameters of interest inside the biofilm (Fig. 2e). Local parameters are quantified for each cube and therefore have a spatial and potentially a temporal dependence. Each cube’s location is given by its centre coordinate (x,y,z), which can be expressed as the distance to the biofilm centre, distance to the biofilm surface, or distance to the substratum. Based on the dissection of the biofilm volume into cubes, numerous structural, textural, and fluorescence properties can be calculated for each cube. For example, cube-internal structural parameters are computed, such as local density and local porosity. Structural parameters also include local biofilm surface properties such as biofilm thickness, surface area, surface roughness coefficient, and surface per substrate area, which are calculated per pillar of cubes sharing the same substrate area, and we assign the same value of these parameters to each cube in a given pillar. Textural parameters are quantified according to Haralick *et al*.^22^, for each cube. Fluorescence parameters calculated for each cube are basic fluorescence quantifications for each fluorescence channel, as well as their overlap, their ratio, the Manders’ overlap and Pearsons’ correlation for different fluorescence channels, autocorrelation, and also density correlation values depending on a range chosen by the user. It is also possible to track pseudo-cell cube lineages (see Supplementary Note) to measure clonal cluster sizes and similar properties.

In addition to the local parameters described above, BiofilmQ also calculates global parameters for the whole biofilm (Fig. 2e). Some of these parameters characterize the size and morphology of the whole biofilm, including its volume, mean thickness, surface area, roughness coefficient, or area covered on the substratum, as well as several combinations of these values, such as the surface-to-volume ratio. A subset of these parameters is also available in COMSTAT^5^, for which we chose identical implementation to enable compatibility (Supplementary Table 1, Supplementary Note). In addition to these structural parameters, BiofilmQ offers the possibility to quantify correlations between different fluorescence reporters globally and locally *via* the Manders’ overlap coefficients, Pearson’s correlation coefficient, volume overlap fractions, and relative abundances of biomass. These parameters enable, *e.g*., quantitative measures of species cluster sizes and separation distances in multispecies biofilms *via* 3D correlation functions. Furthermore, the mean (optionally weighted by cube biovolume to account for cubes with different biovolume fraction), median, minimal, maximal as well as percentiles of every local parameter also make up one global parameter each, which are all calculated for the entire biofilm, its core (cubes deep below the biofilm surface), and shell (cubes close to the surface).

After the analysis of a single 3D (or 2D) biofilm image, BiofilmQ can apply the same analysis to a whole time-series (to analyse the temporal variation of a single biofilm), or to a non-time-series collection of biofilm images (to analyse the variation within a population of biofilms). All data analysis operations can be performed in high-throughput through BiofilmQ-inbuilt batch-processing capabilities, and results can be exported to standard formats (Fig. 2f) or directly visualized (Fig. 2g).

After the image analysis, BiofilmQ offers flexible and powerful options to visualize all quantified parameters (Fig. 3), resulting in publication-quality editable figures. In a spatiotemporal kymograph, the spatial dependence of a biofilm-internal property (*e.g*. a fluorescent reporter or any other cube parameter) can be visualized over time (Fig. 3b). Importantly, different biofilm-internal spatial measures, such as distance-to-surface or distance-to-substrate, can be chosen on the y-axis for these heatmaps, and different temporal measures, such as biofilm volume or surface area, can be also used instead of time on the x-axis. Each column of such kymographs can be plotted as a “1.5D histogram”, in which, for a particular time-point, the biofilm cube parameter of interest can be averaged across cubes within a biofilm and plotted with error bars against a spatial axis of choice (Fig. 3b, inset). If instead of a time series, a population of different biofilms is analysed, choosing the kymograph-plot type will result in a demograph that visualizes variations of the internal spatial structure or fluorescence properties across different biofilms in the population (Fig. 3b)

**Figure 3:**
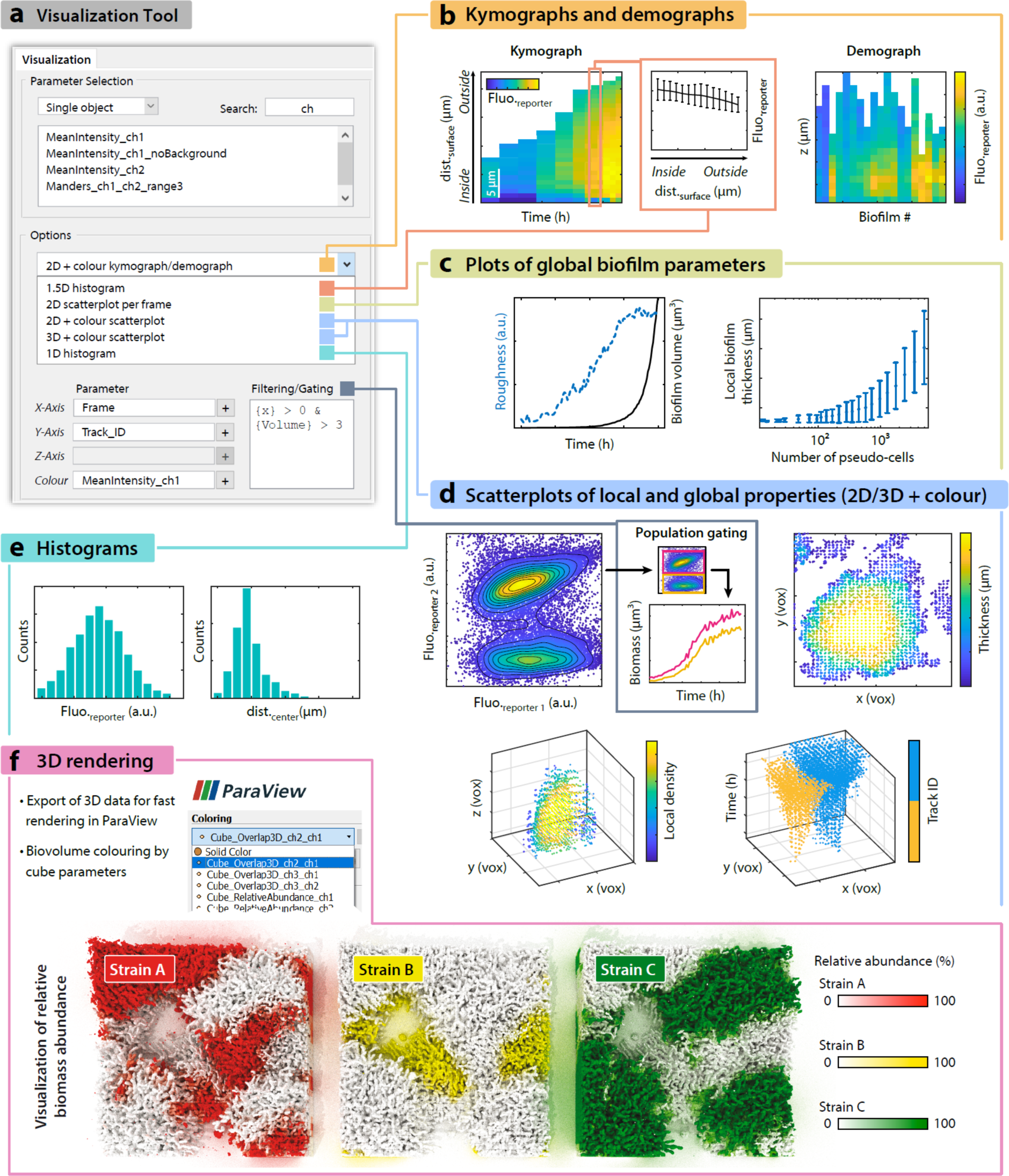
Overview of the BiofilmQ visualization process and capabilities. **a**, Screenshot showing several key elements of the graphical user interface for visualization, depicting how to choose the axis of figures to be plotted, and the plot type. **b**, Left, a kymograph quantifies fluorescent reporter expression as a space-time heatmap. In this case, the fluorescence of an RpoS-mRuby3 translational fusion is plotted over time and space during *V. cholerae* C6706 wild type (WT) biofilm development. Centre, a 1.5D histogram reveals the relation between fluorescent reporter intensity and position in the biofilm for a single time-point, error bars represent the SD across cubes. Right, the heatmap represents a demograph, which reveals spatially-resolved differences between biofilms, for a particular cube-level parameter (here: RpoS-mRuby3 fluorescence as a function of height in *V. cholerae* biofilms after 18 h of growth). **c**, Left, several global biofilm parameters can be plotted into the same figure for better comparison, in this case the biofilm roughness and volume during biofilm development of a *V. cholerae* N16961 rugose Δ*crvA* strain. Right, to observe the behaviour of a parameter in a time series, other time-related quantities, such as the number of pseudo-cell cubes, can be used as timescale on the x-axis. **d**, Top left, analogous to flow cytometry, BiofilmQ can perform biofilm image cytometry, comparing two fluorescent reporters or any other pseudo-cell cube parameters. Here, results are shown from a biofilm co-culture of two *V. cholerae* N16961 WT strains that constitutively produce sfGFP; one of these strains additionally produces mRuby2 constitutively. The segmentation was performed on the sfGFP channel. The gating/filtering option enables the separation of two populations; properties of each gated population can then be visualized separately. Top right, in a 2D+colour scatter plot, extracted cube parameters can be visualized, in this case for example the biofilm thickness distribution in space at a specific timepoint (22 h) during *V. cholerae* C6706 WT biofilm development. Bottom left, similarly, a 3D+colour scatterplot visualizes quantified cube parameters, but provides one additional axis. In this case, the spatial distribution of the local density during *V. cholerae* C6706 WT biofilm growth is shown at a particular timepoint (12 h). Bottom right, visualization of two tracked *V. cholerae* N16961 rugose Δ*crvA* biofilm colonies with the same constitutive fluorescent protein expression (sfGFP), growing together over time. Separation of lineages is performed *via* cube tracking. **e**, Histograms of quantified cube parameters, in this case the fluorescence signal of an RpoS-mRuby3 translational fusion reporter (left) and the distance of each cube to the biofilm centre (right), for a *V. cholerae* C6706 WT biofilm grown for 22 h. Such histograms can help to set thresholds for gating or to better understand the biofilm structure. **f**, Exported files can be opened and edited with other software for further visualization. In this case, the location and globally-computed relative abundance of three different *V. cholerae* N16961 rugose strains (differing by the colour of a constitutively expressed fluorescent protein marker: mTFP1, mKOκ, or mKate2) were calculated using BiofilmQ and the result was exported and rendered in 3D with the ParaView software.

To visualize a global property as a function of time (Fig. 3c), or any other parameter, a simple “2D scatterplot per frame” may be used. The “2D+colour scatterplot” option does not perform averaging per frame and can therefore be used with the axes chosen more freely, for example including spatial coordinates, and can also colour-code each data point according to another parameter (Fig. 3d). Even more flexibility is provided by the “3D+colour scatterplot” for visualizing the quantified local properties of biofilms (Fig. 3d).

Analogous to flow cytometry, the biofilm image cytometry provided by BiofilmQ allows users to apply gates/filters to their data for each pseudo-cell cube, to effectively select pseudo-cell sub-populations with certain characteristics (Fig. 3d, inset). The different gated populations can then be analysed separately with the BiofilmQ plotting capabilities.

To generate 3D rendered biofilms, for which each cube-parameter can be mapped as colour onto the rendered biomass (Fig. 3f), BiofilmQ can export the image analysis results into vtk-files, which can be loaded into the open-source 3D-rendering software ParaView^23^.

BiofilmQ closes a critical gap in the toolset for the spatial and spatiotemporal analysis of microbial communities, by providing a comprehensive image analysis platform that is capable of handling any community spatial geometry. BiofilmQ is user-friendly and designed such that it can be used by beginners within minutes, owing to the extensive video tutorial support and technical documentation. Previously inaccessible global and local biofilm parameters can now be extracted to quantify phenotypic differences between strains, species, and growth conditions, enabling the detailed exploration of phenotypic, genotypic, and structural heterogeneity within biofilms and between biofilms. Batch-processing capabilities make BiofilmQ suitable for biofilm phenotyping in high-content screens. BiofilmQ therefore enables new directions for biofilm research, and simultaneously provides a solid quantitative foundation for future studies of spatially structured microbial communities.

## Methods

### Bacterial biofilm growth

*Vibrio cholerae, Bacillus subtilis*, and *Escherichia coli* strains were routinely grown in liquid lysogeny broth (LB-Miller) at 37 °C under shaking conditions. The strains and plasmids used in this study are listed in Supplementary Tables 2 and 3.

Flow chamber biofilm experiments were performed in M9 minimal medium with 0.5% (w/v) glucose for *V. cholerae*, and tryptone broth (10 g L^−1^) for *E. coli*. To grow flow chamber biofilms of *V. cholerae* and *E. coli*, microfluidic chambers of 7 mm length and 500 µm × 100 µm cross-section were used^18,21^, and a flow rate of 0.1 µL min^−1^ was set using a syringe pump (Pico Plus, Harvard Apparatus). To inoculate flow chambers, overnight cultures were back-diluted 1:200 in LB medium for *V. cholerae*, and 1:80 in 0.9% (w/v) NaCl for *E. coli*, and grown to an optical density at 600 nm of OD_600_ = 0.5. This culture was then used to inoculate the flow chambers. After inoculation, cells were given 1 h to attach to the surface before the constant flow with fresh medium was initiated.

To grow mixed biofilms of *V. cholerae* strains containing different fluorescent protein markers, cultures of all strains were inoculated in microfluidic chambers in a 1:1 ratio for 2 strains, and 1:1:1 for 3 strains, before the constant flow of fresh medium was started.

Pellicle biofilms of *B. subtilis* NCBI3610 carrying P*_tapA_*-*gfp* and P*_tapA_*-*mKate* transcriptional reporters on the chromosome, were grown in MSgg medium^24^ without shaking at the air-liquid interface in 24-well microtiter plates for 48 h at 30°C.

Macrocolony biofilms of *E. coli* AR3110 were initiated by spotting 5 µL of overnight culture on solid LB medium (1.5% agar, w/v). Plates were sealed with parafilm and incubated for 5 days at 23°C, prior to imaging.

### Imaging

For spatiotemporal measurements of different reporters and for separating different populations in flow chambers, biofilms were imaged with a Yokogawa CSU confocal spinning disk unit mounted on a Nikon Ti-E inverted microscope using a Plan Apo 60x oil NA 1.4 objective (Nikon), by exciting fluorescence with a 488 nm laser (for sfGFP) and a 552 nm laser (mRuby2/mRuby3). Images were acquired with an Andor iXon EMCCD camera, cooled to – 80°C, using the EM-gain of the camera. Macrocolony biofilms of *E. coli* strain (KDE1469) were imaged using the above microscope setup, but with a 4x air NA 0.2 objective, exciting the constitutively expressed sfGFP.

To image a mixed population of strains with different fluorescent reporters, and to image pellicle biofilms, images were captured using a Zeiss LSM 880 point-scanning confocal laser scanning microscope, with a 40x NA 1.2 water objective for *V. cholerae* and *B. subtilis*. For *E. coli* biofilms, the same microscope was used with a 63x NA 1.4 oil objective.

### Image analysis

As BiofilmQ is open source software, written in Matlab (MathWorks), it is possible to adapt BiofilmQ to particular user requirements. Algorithms used for biofilm pre-processing, segmentation, parameter quantification, and data visualization are described in detail with examples in the online documentation at https://drescherlab.org/data/biofilmQ. All code is freely available, revealing the exact implementation of each data analysis step.

## Acknowledgements

We are grateful to Takuya Ohmura, Konstantin Neuhaus, Daniel Rode, and the Drescher lab for comments and help with trouble-shooting BiofilmQ, as well as Ana L. Gallego Hernandez, William DePas, Jin Hwan Park, Jennifer Teschler, and Dianne K. Newman for image data. This work was supported by grants to K.D. from the Max Planck Society, European Research Council (StG-716734), Human Frontier Science Program (CDA00084/2015-C), Deutsche Forschungsgemeinschaft (SFB 987), the Behrens-Weise-Stiftung, and the Minna-James-Heineman-Stiftung.

## Author contributions

R.H., H.J., E.J., K.D. conceived and designed the project and the graphical user interface. R.H. developed the algorithms, H.J., E.J. extended algorithms. R.H., H.J., E.J. developed the image analysis and data visualization workflows. H.J., E.J., M.B., S.V. created video tutorials. E.J. wrote the documentation. All authors contributed data, analysis ideas, and important discussions. R.H., H.J., E.J., K.D. wrote the paper and created the figures.

## Additional information

Supplementary information is available in the online version of the paper. Video tutorials, a detailed documentation, and downloads are available at https://drescherlab.org/data/biofilmQ.

## Code and data availability

Image data and processed data used in this study are available from the corresponding author upon reasonable request. Software code is available at https://drescherlab.org/data/biofilmQ.

## References

1. Flemming, H.-C. & Wuertz, S. Bacteria and archaea on Earth and their abundance in biofilms. Nat. Rev. Microbiol. 17, 247–260 (2019).

2. Koo, H., Allan, R. N., Howlin, R. P., Stoodley, P. & Hall-Stoodley, L. Targeting microbial biofilms: current and prospective therapeutic strategies. Nat. Rev. Microbiol. 15, 740–755 (2017).

3. Stewart, P. S. & Franklin, M. J. Physiological heterogeneity in biofilms. Nat. Rev. Microbiol. 6, 199–210 (2008).

4. Nadell, C. D., Drescher, K. & Foster, K. R. Spatial structure, cooperation and competition in biofilms. Nat. Rev. Microbiol. 14, 589–600 (2016).

5. Heydorn, A. et al. Quantification of biofilm structures by the novel computer program COMSTAT. Microbiology 146 (Pt 10), 2395–407 (2000).

6. Daims, H., Lücker, S. & Wagner, M. daime, a novel image analysis program for microbial ecology and biofilm research. Environ. Microbiol. 8, 200–13 (2006).

7. Mueller, L. N., de Brouwer, J. F. C., Almeida, J. S., Stal, L. J. & Xavier, J. B. Analysis of a marine phototrophic biofilm by confocal laser scanning microscopy using the new image quantification software PHLIP. BMC Ecol. 6, 1 (2006).

8. Chávez de Paz, L. E. Image analysis software based on color segmentation for characterization of viability and physiological activity of biofilms. Appl. Environ. Microbiol. 75, 1734–9 (2009).

9. Meijering, E., Carpenter, A. E., Peng, H., Hamprecht, F. A. & Olivo-Marin, J.-C. Imagining the future of bioimage analysis. Nat. Biotechnol. 34, 1250–1255 (2016).

10. Sliusarenko, O., Heinritz, J., Emonet, T. & Jacobs-Wagner, C. High-throughput, subpixel precision analysis of bacterial morphogenesis and intracellular spatio-temporal dynamics. Mol. Microbiol. 80, 612–627 (2011).

11. Ducret, A., Quardokus, E. M. & Brun, Y. V. MicrobeJ, a tool for high throughput bacterial cell detection and quantitative analysis. Nat. Microbiol. 1, 16077 (2016).

12. Paintdakhi, A. et al. Oufti: an integrated software package for high-accuracy, high-throughput quantitative microscopy analysis. Mol. Microbiol. 99, 767–77 (2016).

13. Hartmann, R., Teeseling, M. C. F. van, Thanbichler, M. & Drescher, K. BacStalk: a comprehensive and interactive image analysis software tool for bacterial cell biology. bioRxiv 360230 (2018). doi:10.1101/360230

14. Linkert, M. et al. Metadata matters: access to image data in the real world. J. Cell Biol. 189, 777–82 (2010).

15. Yang, X., Beyenal, H., Harkin, G. & Lewandowski, Z. Evaluation of biofilm image thresholding methods. Water Res. 35, 1149–58 (2001).

16. Yerly, J., Hu, Y., Jones, S. M. & Martinuzzi, R. J. A two-step procedure for automatic and accurate segmentation of volumetric CLSM biofilm images. J. Microbiol. Methods 70, 424–33 (2007).

17. Renslow, R., Lewandowski, Z. & Beyenal, H. Biofilm image reconstruction for assessing structural parameters. Biotechnol. Bioeng. 108, 1383–94 (2011).

18. Drescher, K. et al. Architectural transitions in Vibrio cholerae biofilms at single-cell resolution. Proc. Natl. Acad. Sci. U. S. A. 113, E2066–E2072 (2016).

19. Wang, J. et al. Bact-3D: A level set segmentation approach for dense multi-layered 3D bacterial biofilms. in 2017 IEEE International Conference on Image Processing (ICIP) 330–334 (IEEE, 2017). doi:10.1109/ICIP.2017.8296297

20. Luo, T. L. et al. A Sensitive Thresholding Method for Confocal Laser Scanning Microscope Image Stacks of Microbial Biofilms. Sci. Rep. 8, 13013 (2018).

21. Hartmann, R. et al. Emergence of three-dimensional order and structure in growing biofilms. Nat. Phys. 15, 251–256 (2019).

22. Haralick, R. M., Shanmugam, K. & Dinstein, I. Textural Features for Image Classification. IEEE Trans. Syst. Man. Cybern. SMC-3, 610–621 (1973).

23. Ahrens, J., Geveci, B. & Law, C. ParaView: An End-User Tool for Large-Data Visualization. in The Visualization Handbook (eds. Johnson, C. R. & Hansen, C. D.) 717–731 (Elsevier, 2005).

24. Dragoš, A. et al. Division of Labor during Biofilm Matrix Production. Curr. Biol. 28, 1903–1913.e5 (2018).

